# A Metagenomic Approach Does Not Elucidate Uremic Toxin Levels in Hemodialysis Patients

**DOI:** 10.1101/2024.07.16.603811

**Authors:** Diana H. Taft, Julia M. Quinones, Melissa L. Moreno, Asmaa Fatani, Joonhyuk Suh, Gabrielle Gorwitz, Yu Wang, Mark S. Segal, Wendy J. Dahl

## Abstract

Blood levels of uremic molecules generated through gut microbial metabolism are associated with disease risk, reduced quality of life, and mortality in patients with renal insufficiency. Modulation of the microbiome may offer therapeutic potential.

**Objective:** This study aimed to elucidate the relationships between fecal microbiome and serum levels of targeted uremic molecules in adults undergoing hemodialysis and, secondarily, the role of relative macronutrient substrate availability.

**Methods:** Fecal microbiota was profiled by whole metagenome sequencing, serum *p*-cresyl sulfate (CS), indoxyl sulfate (IS), phenylacetylglutamine (PAG), and trimethylamine *N*-oxide (TMAO) were quantified by liquid chromatography–tandem mass spectrometry (LC-MS/MS), and dietary intake was assessed by three multipass-method 24-hr recalls.

**Results:** Differences in gut microbiome associated with serum uremic toxin levels were not detected. Instead, the relative substrate availability for the microbiota, using dietary protein-to-fiber ratio, was significantly associated with uremia. Serum levels of IS (multivariate linear model, p=0.042) and TMAO (p=0.032) were positively associated with dietary protein-to-fiber ratio, but not CS (p=0.096) and PAG (p=0.44).

**Conclusion:** The lack of association of fecal microbiome with serum uremic toxins suggests that hemodialysis patients possess sufficient microbial enzymatic capacity for the synthesis of these molecules and that, instead, microbially available substrate, protein vs. fiber, may be the primary driver of production.

## Introduction

Diminishing kidney function is the major contributing factor to circulating uremic toxins – the inflammatory microbial-generated metabolites that contribute to significant morbidity^1^ and reduced quality of life^2^ in the chronic kidney disease (CKD) population. The highest circulating levels of protein-bound uremic toxins are typically seen in those with end-stage renal disease (ESRD),^3^ as hemodialysis offers limited removal. However, blood levels of these compounds demonstrate high interindividual variation among hemodialysis patients.^4,5^ suggesting that other factors, such as diet and microbiota composition, influence uremic status.

Logically, macronutrient intake provides varied substrates for microbial fermentation, saccharolytic vs. proteolytic, which would be expected to influence uremic toxin production. Evidence suggests that the dietary protein-to-fiber ratio is correlated with blood levels of uremic toxins.^6,7^ Higher protein intake provides precursor aromatic amino acids for the microbial synthesis of such toxins as *p*-cresyl sulfate, indoxyl sulfate, and phenylacetylglutamine, increasing levels, and conversely, higher fiber intake, provides microbial-available carbohydrates that suppress proteolytic activity through various mechanisms.^8,9^ However, from a clinical patient perspective, mitigation of uremic toxin production through diet modification offers considerable challenges. Limiting dietary protein is not a feasible therapeutic option, given the high prevalence of protein-energy malnutrition in the hemodialysis patient population ^10^ and reported protein intakes typically fall far below recommendations.^11,12^ Alternatively, higher fiber diets could be recommended to favorably shift the protein-to-fiber ratio. To date, the limited clinical trials of prebiotic fiber supplementation have shown mixed results.^11,13-16^ A vegetarian dietary pattern providing a diverse array of dietary fibers shows promise for mitigating uremic toxin blood levels^17^ but may be particularly challenging to implement in the general hemodialysis patient population because of socioeconomic factors.^15^

Gut microbiota composition by 16S rRNA-based sequencing of hemodialysis patients compared to healthy controls^18-21^ and earlier stages of CKD^22^ suggest dysbiosis but with conflicting outcomes as to the specific disruptions. Some studies report suppression of various butyrate-producing bacteria^18,22^ and health-enhancing genera such as *Bifidobacterium*.^20^ ESRD patients, specifically, have been shown to exhibit higher levels of *Klebsiella* and lower levels of *Roseburia* and *Blautia* compared to healthy controls.^23^ Similarly, Wang et al. reported depleted *Roseburia* but also lower *Prevotella, Clostridium, Faecalibacterium*, and *Eubacterium rectale*, and higher levels of *Eggerthella, Flavonifractor, Alistipes, Ruminococcus*, and *Fusobacterium spp*.^24^ Additionally, microbial diversity and its genetic capacity differ in ESRD compared to healthy controls.^24^ Within the hemodialysis population, the effects of obesity^25^ and various therapeutic modalities^20,26^ on microbiota composition have been examined. However, how gut microbiota composition and activity contribute to uremic toxin generation has been underexplored. Variations in uremic toxin concentrations of hemodialysis patients consuming similar poor-quality diets^27^ suggest that there may be differences in microbial production (and subsequent absorption) related to microbiota composition and its metabolic activity, which may not be directly or completely driven by substrate availability.

Manipulation of the microbiome offers a novel potential therapy to mitigate uremic toxin production and, thus, subsequent absorption of these metabolites. Given the evidence for gut microbiota dysbiosis of hemodialysis patients,^28^ achieving clinically relevant reductions in uremic toxins may require intentional targeting of the responsible microbial species or of the responsible microbial pathways in addition to implementing healthful dietary approaches. Therefore, the aim of this analysis was to elucidate the relationships between fecal microbiome composition and function using in-depth whole metagenomic sequencing of CKD patient feces and serum levels of targeted uremic molecules in adults undergoing hemodialysis. A secondary aim was to determine if a proxy for relative substrate availability, defined as the dietary protein-to-fiber ratio, was associated with serum uremic levels.

## Methods

Fecal samples from hemodialysis patients participating in two studies at one dialysis center in <blinded>, USA, were analyzed by whole metagenomic sequencing. As previously reported, baseline fecal samples were collected from adults undergoing hemodialysis and participating in a clinical trial examining the effect of fiber supplementation on uremic molecules and fecal microbiota composition.^11^ A follow-up cross-sectional study included adults receiving hemodialysis, and those with known intestinal disease (i.e., inflammatory bowel disease, malignancy, celiac disease) or previous colorectal surgery were excluded. Stool collections were made using Fisherbrand™ Commode Specimen Collection System (Fisher Scientific, cat. no. 02-544-208), kept on ice for up to 6 h until processing, and stored at -80°C for later DNA extraction. The study protocols were approved by the Institutional Review Board of <blinded>, and the clinical trial was registered at clinicaltrials.gov <blinded>. Written informed consent was obtained from all participants of the original studies, and study procedures were in accordance with the Declaration of Helsinki.

### Calculation of Dietary Protein to Fiber Ratio

As a proxy of microbial substrate availability, dietary protein and fiber intake, was assessed using a repeated, multipass 24-hour recall method.^29^ As previously described, nutrient intake was estimated using Food Processor Nutrition Analysis Software (ESHA version 11.3.2).^11^ Mean protein (g per day) and fiber (g per day) over 3 non-consecutive days were determined, and the ratio of protein to fiber intake was calculated for each study participant.

### Quantification of Uremic Metabolites

The predominate microbial-derived uremic toxins of the serum metabolome in ESRD, indoxyl sulfate (IS), *p*-cresyl sulfate (CS), trimethylamine-*N*-oxide (TMAO), phenylacetylglutamine (PAG) were targeted.^24^ Pre-dialysis blood was collected via hemodialysis vascular access following one day of non-dialysis. Methods for the quantification of serum levels of uremic molecules by liquid chromatography–tandem mass spectrometry (LC-MS/MS) have been previously described.^11^

### DNA Extraction and Sequencing

DNA from feces was extracted using the Qiagen AllPrep PowerFecal DNA/RNA kit (cat no 80244) per manufacturer instructions. DNA was quantified using the QuBit dsDNA BR Assay kit (cat no Q32850) per the manufacturer’s instructions. After confirming that samples were of adequate quantity, samples were submitted to the UF ICBR NextGen DNA sequencing core for library preparation using Illumina Standard Library Construction (New England Biolabs). DNA sequencing was completed by the UF ICBR using Illumina NovaSeq S4 150 PE chemistry.

### Bioinformatic Processing and Statistical Analysis

Host subtraction was completed using BMTagger,^30^ and trimming was completed using Trimmomatic.^31^ Functional assignment was completed using HUMAnN3.^32^ Metaphlan3 was used for taxonomic assignment.^32^ Additional analysis was completed using R version 4.2.2. The distance measure used for beta-diversity analysis was the robust Aitchison’s distance as implemented in the robCompositions package.^33^ Beta diversity differences by species were visualized using NMDS based on robust Aitchison’s distance in the vegan package.^34^ Differences in species beta diversity by uremic toxin molecules were tested using PERMANOVA as implemented in the adonis2 function in the vegan package. Beta diversity based on gene abundance was visualized in the same way as species abundance (again, based on the robust Aitchison’s distance), and adonis2 was again used to test for differences in gene abundance. Maaslin2 was used to test for significant differences in species and individual gene abundance.^35^ The relationship between uremic toxin molecules and dietary protein-to-fiber ratio was tested with a multivariate linear model using the lm function in R, with the uremic toxin molecules as the outcome and the protein-to-fiber ratio as the primary predictor. Additional covariates, patients’ sex and age, were tested for possible inclusion using ANOVA (the ANOVA function in R) to compare the model with and without these terms to determine if the model improved with their inclusion.

## Results

A total of 21 participants had baseline stool samples remaining from the original studies for inclusion in this study. Most participants were African American and on dialysis for more than 2 years. Table 1 has additional demographic information.

Regarding the composition of the fecal microbiome of all participants, the most abundant taxa at the family level was Bacteroidaceae, with a median relative abundance of 28.6% (range 0.2% to 58.3%). Other taxa at the family level with a median relative abundance greater than 1% included Coriobacteriaceae with a median relative abundance of 1.4% (range 0.0% to 13.5%), Eubacteriaceae (1.7%; range 0.1% to 13.6%), Lachnospiraceae (24.1%; range 11.1% to 70.0%), Rikenellaceae (1.9%; range 0.0% to 13.3%), Ruminococcaceae (13.3%; range 0.9% to 40.7%), and Tannerellaceae (1.9%; range 0.0% to 6.7%) (Figure 1). Examination of the genes present across all samples showed UniRef90_A7V4G2 (uncharacterized predicted protein from *Bacteroides uniformis* whole genome sequencing data without similarity to characterized proteins at the 90% identity level) with a median abundance of 1228 (range 0 to 3297) to be the most abundant gene. Other abundant genes, all uncharacterized proteins without similarity to characterized proteins at the 90% identity level, were UniRef90_A0A0E1X896 with a median abundance of 558.8 (range 4.4 to 1284.6), UniRef90_A0A395VRV3 (600.3; range 0.0 to 1877.3), UniRef90_A5ZYV3 (596.4; range 0.0 to 3058.8), UniRef90_A7B5X2 (367.0; range 185.7 to 844.6), and UniRef90_E2N7L5 (393.2; range 0.0 to 1412.5).

**Figure 1.**
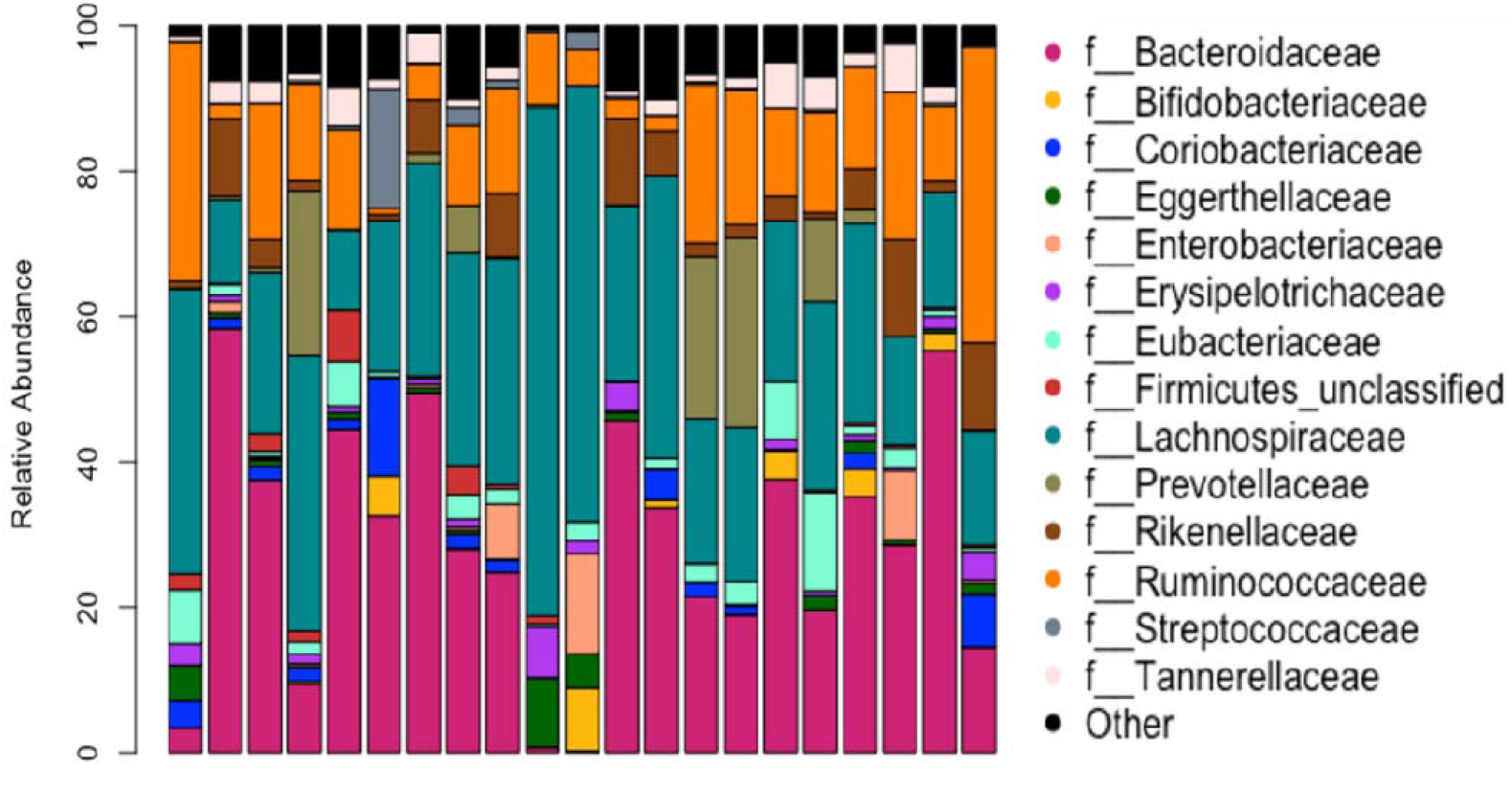
Relative abundance of taxa at the family level. Each bar represents a fecal sample from a single participant.

Following the characterization of the taxa and genes present in all samples, we sought to determine if there were any differences in either taxa or gene abundance by blood levels of uremic toxins using MaAsLin2. No significant differences in either taxa or gene abundance associated with levels of any of the measured uremic toxin molecules were found. For beta diversity, there was no apparent clustering of species composition by uremic toxin blood levels for any of the studied uremic toxins (Figure 2). With all uremic toxin levels included in a single model, PERMANOVA results based on Robust Aitchison distance and PERMANOVA confirmed that there were no significant differences in beta diversity by any of the studied uremic toxins (CS, p=0.91; IS, p=0.83; PAG, p=0.41; TMAO, p=0.16).

**Figure 2.**
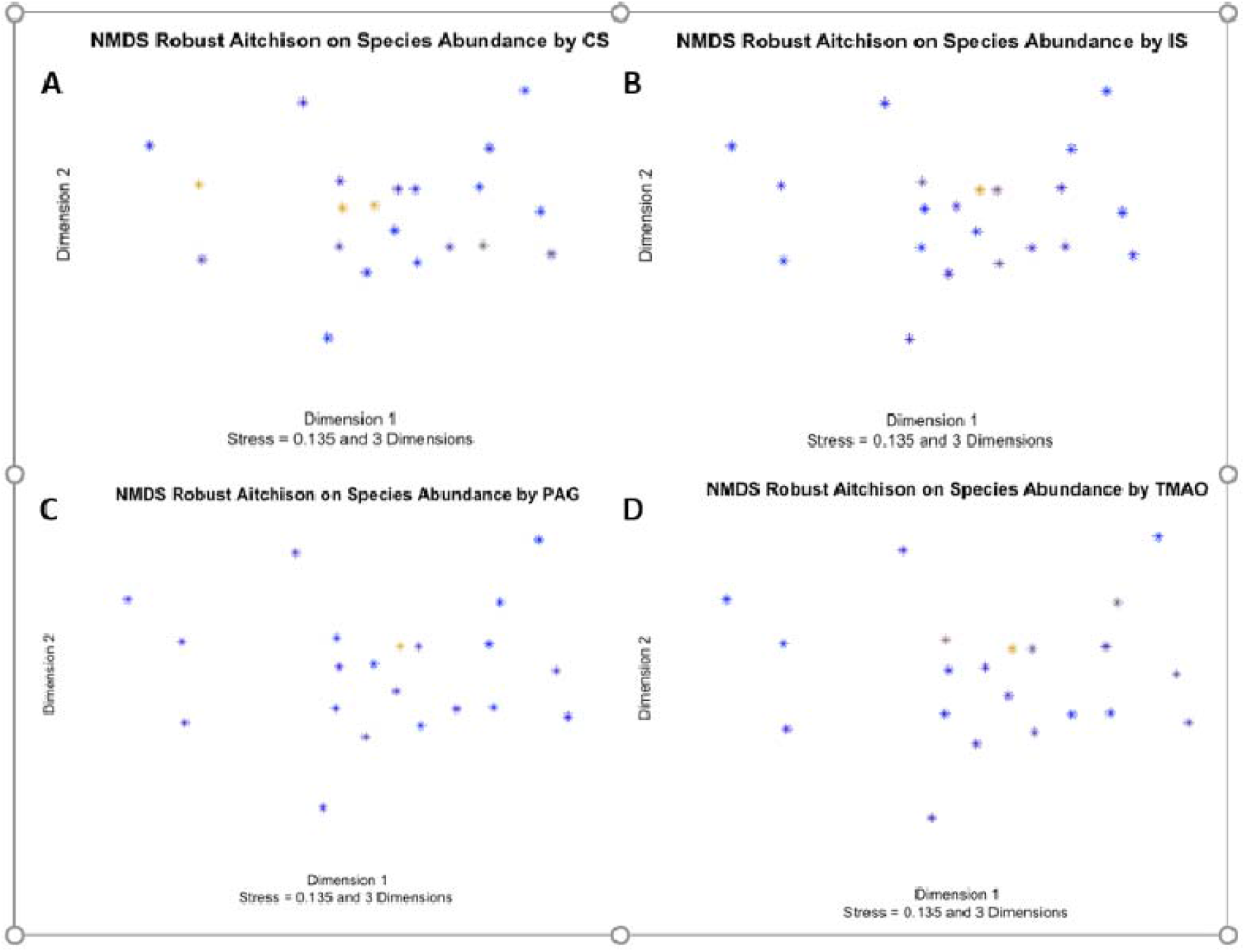
NMDS based on Robust Aitchison distance of species relative abundance. NMDS was run with 3 dimensions, only the first two dimensions are shown in these plots. Blue points indicate low levels of uremic toxin molecules in a gradient to yellow, indicating the highest levels. (A) Plot for *p*-cresyl sulfate (B) Plot for indoxyl sulfate (C) Plot for phenylacetylglutamine (D) Plot for trimethylamine-*N*-oxide.

The gene-abundance-based beta-diversity analysis had results comparable to those for beta diversity based on species. There was no visible separation on NMDS plots by uremic toxin levels (Figure 3) nor significant differences in gene-abundance-based beta diversity by any uremic toxin (CS, p=0.88; IS, p=0.53; PAG, p=0.97; TMAO, p=0.89) based on robust Aitchison distance and PERMANOVA.

**Figure 3.**
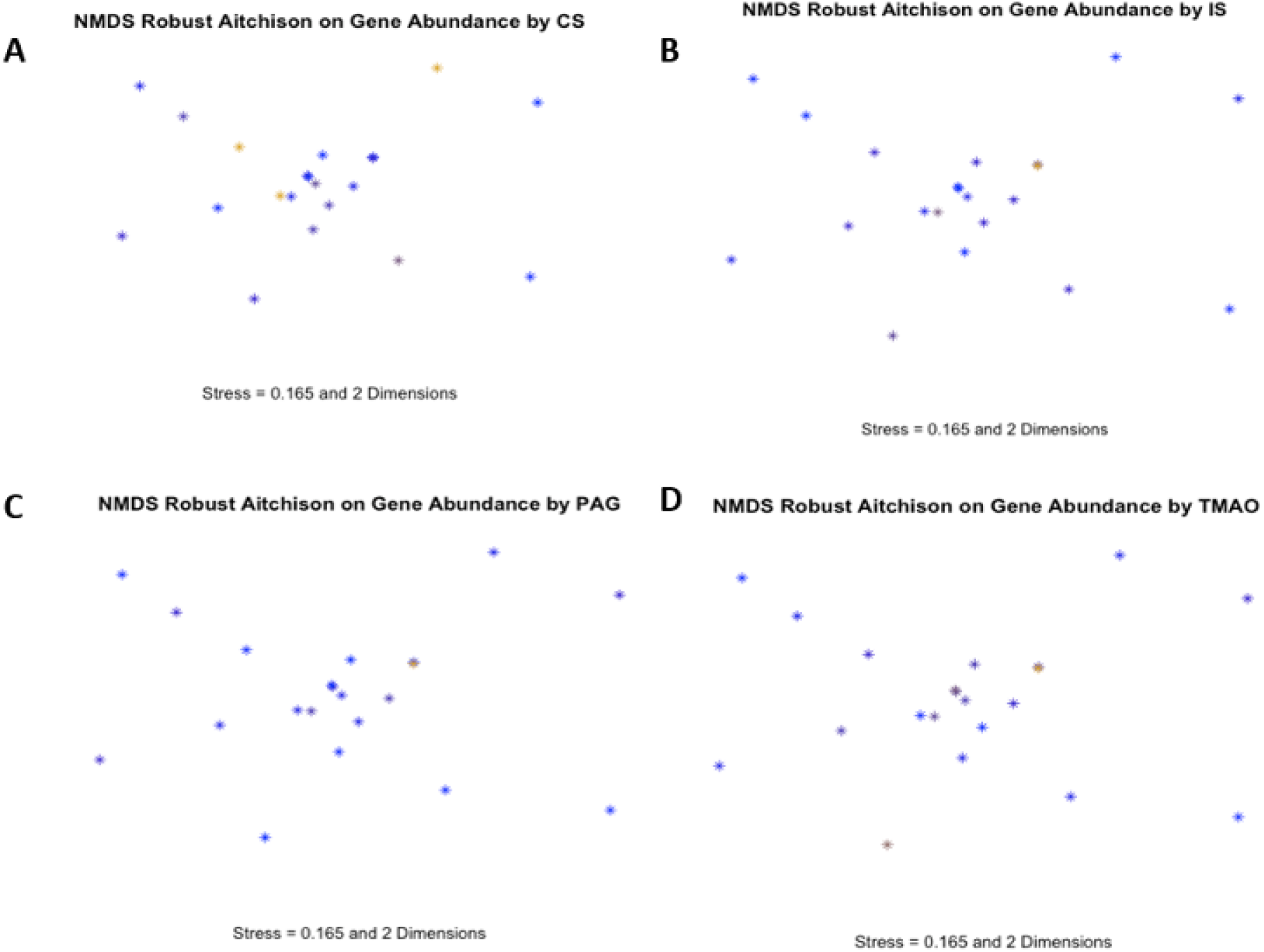
NMDS based on robust Aitchison distance of species relative abundance. NMDS was run with 2 dimensions. Blue points indicate low levels of uremic toxin molecules in a gradient to yellow, indicating the highest levels of uremic toxin molecules. (A) Plot for *p*-cresyl sulfate (B) Plot for indoxyl sulfate (C) Plot for phenylacetylglutamine (D) Plot for trimethylamine-*N*-oxide.

For the test of association between protein-to-fiber ratio and the individual uremic toxins, the inclusion of sex and age did not significantly improve the model (ANOVA, p=0.12). Therefore, protein-to-fiber ratio was the only predictor variable retained in the model. Protein-to-fiber ratio was significantly associated with IS (p=0.042) and TMAO (p=0.032) but not CS or PAG (Table 2). For IS and TMAO, an increased protein-to-fiber ratio (higher protein and lower fiber consumption) was associated with increased uremic toxin levels.

## Discussion

The gut microbiome of ESRD patients presents as dysbiotic.^5,21,24^ Functional capacity suggests depletion of amino acid biosynthesis and enhanced degradation, suggesting increased availability of amino acids^24^ – substrate for the generation of uremic toxins.^36^ We hypothesized that gut microbiome profiles of hemodialysis patients would differ based on the serum level of microbial-generated uremic toxins, a proxy of microbial production. Surprisingly, no associations between gut microbiome and levels of uremic toxins were detected in this study. It is possible that the microbiome may exert an effect in patients with higher residual renal function and that the lack of association may be due to the advanced stage of renal disease and its associated dysbiosis. The finding suggests that the fecal microbiomes of the hemodialysis patients assessed had sufficient levels of relevant microbes and functional pathways required to produce uremic toxins. This is supported by previous research demonstrating that ESRD patients exhibit a higher abundance of genes encoding for uremic toxin synthetases compared to healthy controls.^24^ Although gut microbiota composition may be an important determinant of the serum metabolome of ESRD patients when compared to healthy controls,^24^ this may not be the case when they are compared within the ESRD patient population. The larger driver of variations in uremic toxin levels may instead be substrate availability, sourced from the patients’ diets.

The interplay between dietary intake, gut microbiome, and levels of uremic toxins has been previously demonstrated. For example, higher serum TMAO is associated with increased dietary intake of protein, specifically from animal-sourced foods such as red meat,^37^ which provide carnitine and choline for the microbial synthesis of TMA.^38^ Conversely, a vegetarian diet, providing lower levels of these substrates, is associated with lower blood levels of TMAO.^37^ However, limiting the source substrate for CS, IS, and PAG, synthesized from the aromatic amino acids tyrosine, tryptophan, and phenylalanine, respectively, is more challenging, as total protein intake would need to be limited. In adults with CKD, adherence to a low-protein diet mitigates blood levels of CS and IS,^39^ with only minor modulation of gut microbiota composition.^40^ Given the high protein needs of ESRD patients^41^ and their risk of protein-energy malnutrition,^10^ simply limiting protein is contraindicated. A more comprehensive approach to diet therapy is needed. Serum levels of CS and IS are lower in hemodialysis patients consuming a vegetarian diet,^17^ a dietary pattern typically higher in fiber. Furthermore, there is evidence to support that the relative dietary intake of protein to fiber functions to mitigate microbial production of uremic toxins in ESRD;^6,7^ thus, increasing the intake of dietary fiber is needed for uremic toxin mitigation. The findings of the present study, support a direct association between the dietary protein-to-fiber ratio and serum IS, consistent with prior reports.^6,42^ Not unexpectedly, TMAO was also associated with protein-to-fiber ratio in this patient sample of non-vegetarians. Conversely, dietary protein-to-fiber ratio and serum level of CS trended but were not significantly associated, in contrast with previous reports in CKD patients,^7^ those receiving hemodialysis,^6^ and controlled feeding of healthy adults.^43^ This lack of significant effect of CS, in addition to the limitation of sample size, may be due to the generally low to moderate fiber intakes of the hemodialysis patients, i.e., an adequate level of dietary fiber substrate and perhaps decreased transit time is needed to shift from proteolytic to saccharolytic fermentation.^36^ Research is needed to determine if supplying diverse microbial-available carbohydrates vs. selective prebiotic sources is more effective in reducing uremic molecule generation.

Alternative therapeutic approaches to reducing uremic toxin production have received little attention. First, no known therapeutic research has addressed the impairment of the digestion of protein and/or absorption of peptides and amino acids, which is thought to contribute to malnutrition, inflammation, and atherosclerosis in ESRD patients.^44^ Irrespective of protein intake, this impairment will contribute to microbial-accessible substrate destined for the proteolytic fermentation and synthesis of uremic toxins. Potential mechanisms have been suggested ^44^ and include disruption due to small intestinal bacterial overgrowth^45^ and uremia-induced pancreatic exocrine dysfunction.^46^ Research is needed to elucidate the etiology of compromised protein digestion and/or absorption and if impairment differs among ESRD patients. Additionally, therapeutic interventions that may inadvertently elevate uremic toxins, such as the provision of carnitine supplements for the treatment of anemia,^47^ require reconsideration. Supplemental _L_-carnitine is particularly problematic given its low bioavailability^48^ and, thus, high microbial availability^49^ compared to the higher bioavailability of dietary carnitine within a higher protein, adequate fiber diet.^50^ Furthermore, the dietary intake of precursor substrates of other abundant uremic molecules, such as hippuric acid, is also of interest, given their association with cardiovascular disease and mortality.^51^ Interestingly, source substrate for benzoic acid and eventual hippuric acid synthesis are polyphenols from berries^52,53^ and green tea^54^, foods generally thought to promote health.^55^ However, given the limited removal of hippuric acid with dialysis,^56^ research is needed to determine if dietary intake or restriction confers benefit for ESRD patients.

This study had limitations. The analysis was limited to the four most abundant serum uremic toxins in hemodialysis patients, and serum was measured at only one timepoint. Thus, we did not assess the vector of change for the uremic toxins. The small sample size limits our ability to detect differences in the gut microbiome, especially as whole metagenomic sequencing data can be noisy, with multiple different species contributing to similar functions within the gut microbiome.^57^ We are likely underpowered to detect potentially relevant differences in the species composition of the gut microbiome, but given that functional data are more stable, this is less of a problem for the analysis focused on gene pathways. This study was also limited to a single baseline stool sample from each participant. This means if the gut microbiome is changing dynamically in response to dialysis in ways that affect uremic toxin levels, this study was not designed to be able to detect dynamic differences.

Dysbiosis has been established in many chronic disease states.^58^ In ESRD patients, dysbiosis not only contributes to amino acid degradation and the generation of inflammatory uremic toxins but also reduces the synthesis of generally beneficial short-chain fatty acids,^24^ contributing to significant morbidity and mortality. The findings of this study suggest that metatranscriptomics may be useful in better understanding the role of the gut microbiome in uremic toxin production in the hemodialysis patient population, as gene expression levels may provide more meaningful information than just the potential for gene expression. It also suggests that diet composition, irrespective of gut microbes, may explain the wide differences in circulating levels of uremic toxins seen in hemodialysis patients. Well-designed clinical trials are needed to test the feasibility and efficacy of dietary interventions that significantly increase microbial-available carbohydrates - dietary fibers and resistant starches. Medical nutrition therapy is needed to transition hemodialysis patients to a dietary pattern with a reduced protein-to-fiber ratio, whilst preserving protein intake and, thus, the nutritional status of this vulnerable patient population.

### Practical Application

Previous research has shown that gut microbiota compositions of chronic kidney disease differ from healthy cohorts, and blood levels of toxins may be related to these differences. However, in hemodialysis patients, differences in dietary intake, specifically higher protein and lower dietary fiber intakes, and not microbiome, are associated with higher blood levels of toxins. Hemodialysis patients should be encouraged to consume a dietary pattern providing an abundance of dietary fiber while maintaining recommended protein intake.

## Notes

### Competing Interest Statement

The authors have declared no competing interest.

